# Environmental Dependence of the Structure of the C-terminal Domain of the SARS-CoV-2 Envelope Protein

**DOI:** 10.1101/2020.12.29.424646

**Authors:** Kundlik Gadhave, Ankur Kumar, Prateek Kumar, Shivani K. Kapuganti, Neha Garg, Michele Vendruscolo, Rajanish Giri

## Abstract

The SARS-CoV-2 envelope protein (E) is involved in a broad spectrum of functions in the cycle of the virus, including assembly, budding, envelope formation, and pathogenesis. To enable these activities, E is likely to be capable of changing its conformation depending on environmental cues. To investigate this issue, here we characterised the structural properties of the C-terminal domain of E (E-CTD), which has been reported to interact with host cell membranes. We first studied the conformation of the E-CTD in solution, finding characteristic features of a disordered protein. By contrast, in the presence of large unilamellar vesicles and micelles, which mimic cell membranes, the E-CTD was observed to become structured. The E-CTD was also found to display conformational changes with osmolytes. Furthermore, prolonged incubation of the E-CTD under physiological conditions resulted in amyloid-like fibril formation. Taken together, these findings indicate that the E-CTD can change its conformation depending on its environment, ranging from a disordered state, to a membrane-bound folded state, and an amyloid state. Our results thus provide insight into the structural basis of the role of E in the viral infection process.

**Highlights:**

1. The E-CTD of SARS-CoV-2 is intrinsically disordered in solution
2. The E-CTD folds into an ordered structure in presence of membrane mimetics
3. The E-CTD displays conformational changes in the presence of osmolytes
4. Prolonged incubation of the E-CTD leads to its self-assembly into amyloid-like fibrils

**Graphical Abstract:** Structural heterogeneity of the E-CTD.
The E-CTD shows a disordered secondary structure in an aqueous solution and converts into an ordered structure in the presence of membrane mimetics (neutral and negative lipids) and natural osmolytes (TMAO). Incubation at physiological condition shows typical amyloid-like fibrils. The yellow-colored structure represents a predicted structure of the E-CTD by PEP-FOLD.

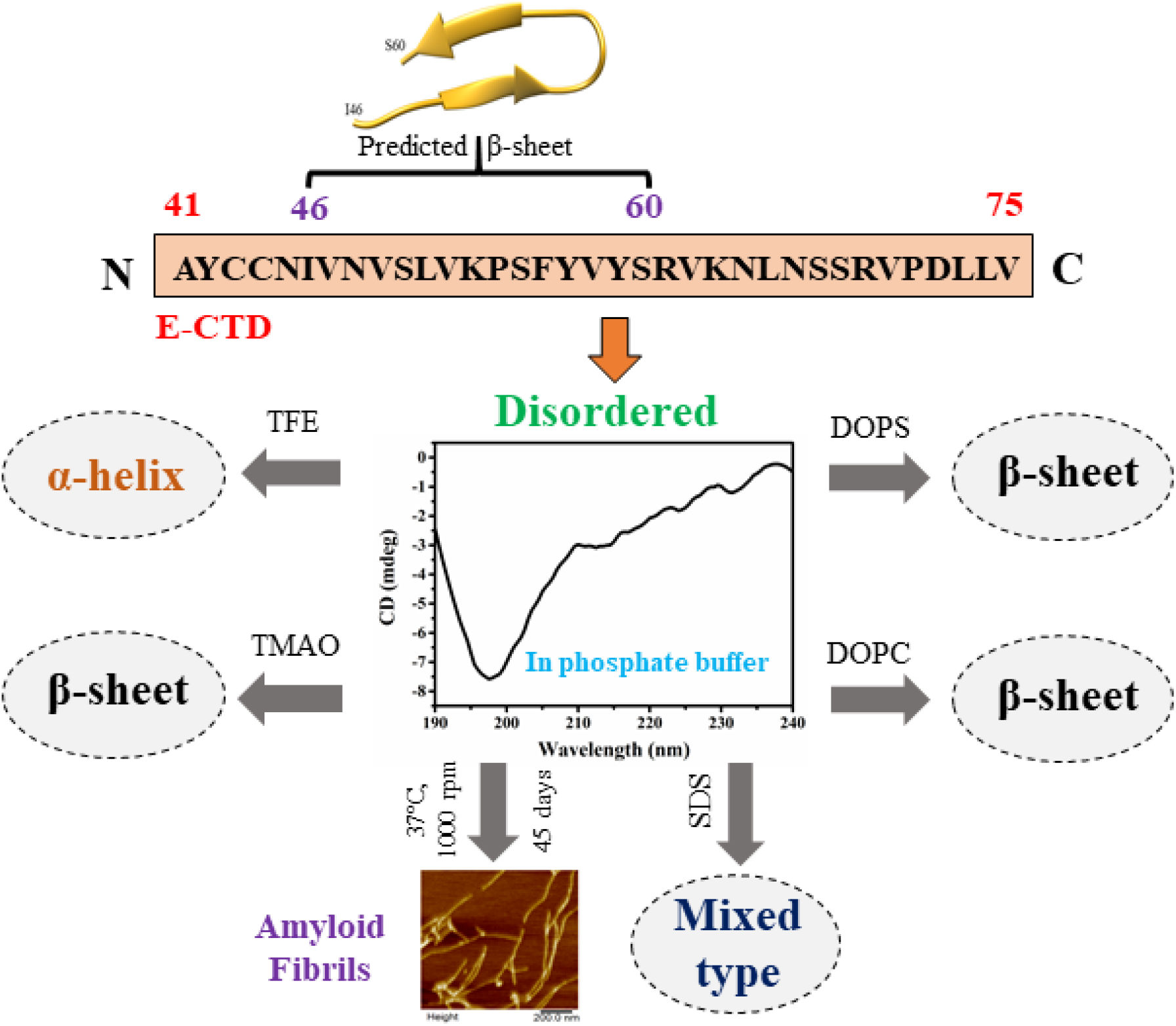

## Introduction

In order to hijack the host cellular machinery for their own replication, viruses are endowed with proteins that mediate the rearrangement of host cell membranes [1,2]. To determine the viral cycle in the host and to develop effective pharmacological treatments, it is therefore important to understand how viral proteins interact with the lipid membranes and proteins of the host cells.

In SARS-CoV-2, it has been shown that the envelope protein (E) can promote changes in the properties of lipid membranes of the secretory organelles of the host cells, which help in packaging and releasing the new viral particles [3,4]. E has thus a dual role, since it acts both as a structural protein, being an essential part of the viral capsid, and as a modulator of the cell membrane properties of the host cell during the replication of the virus. E is a transmembrane protein with a sequence of 76 amino acids, which has 97% sequence similarity with human SARS-CoV (**Figure 1**) and bat SARS-like CoV [4–6]. It consists of a hydrophilic N-terminal peptide (E-NTD), a central hydrophobic transmembrane domain (E-TMD) [7], and a weakly hydrophobic C-terminal domain (E-CTD) [8] (**Figure 1**). In the endoplasmic reticulum (ER) and Golgi of the host cells, the E-NTD is luminally-oriented, the E-TMD has an ion channel activity mediated by the formation of pentameric oligomers [7], and the E-CTD is cytoplasmic-oriented [8]. Through this arrangement, the E-CTD can thus modify the properties of the ER and Golgi membranes, and interact with many host proteins, including Bcl-xL, PALS1, syntenin, sodium/potassium (Na^+^/K^+^) ATPase α-1 subunit, and stomatin [9]. In this way, the E-CTD is crucial for cellular trafficking, in particular by targeting proteins towards Golgi interactions [9].

**Figure 1.**
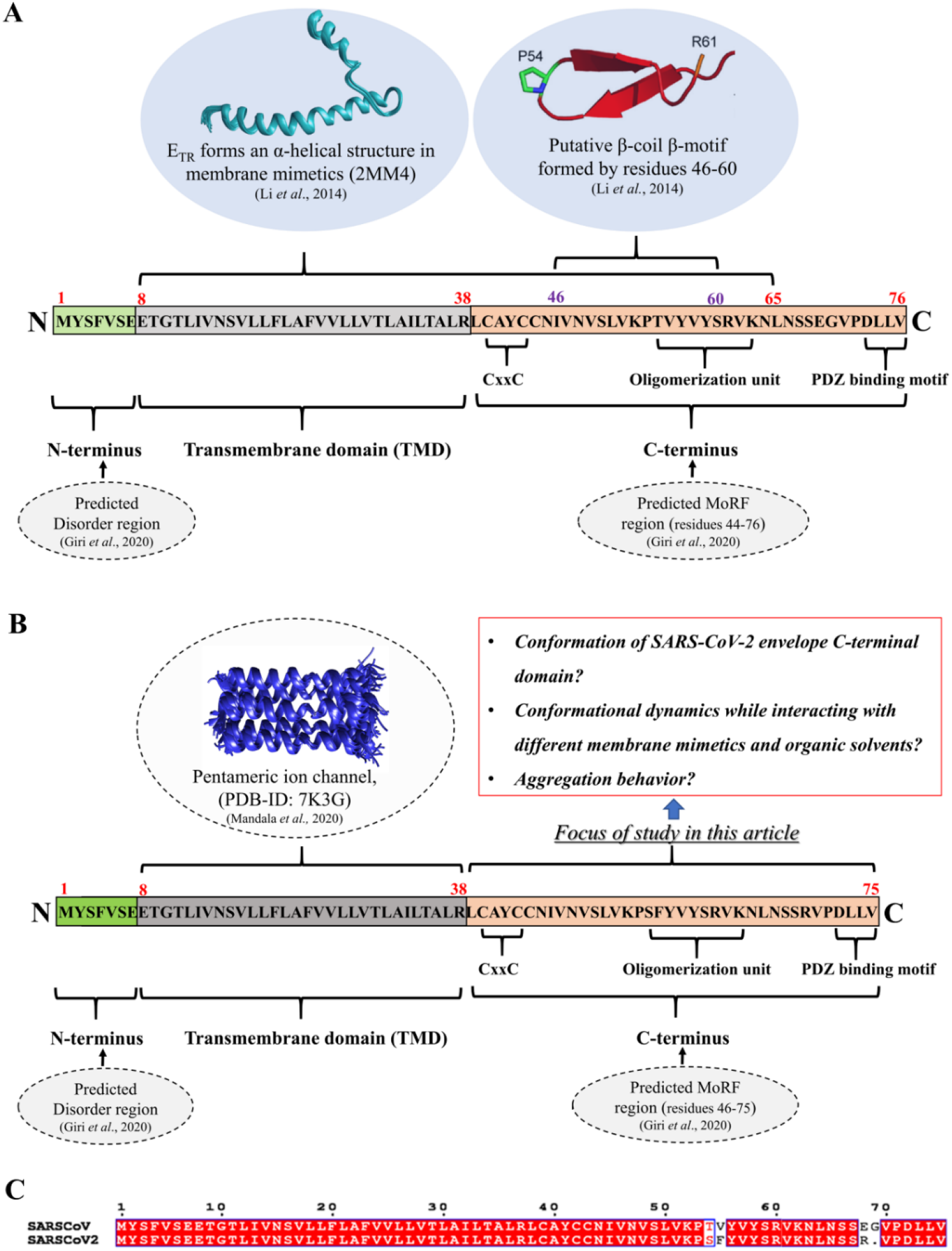
Domain composition of the envelope protein (E) of SARS-CoV and SARS-CoV-2. **(A)** Schematic representation of the reported structural conformations of the three regions in the amino acid sequence of E from SARS-CoV. **(B)** Summary of the reported conformational properties of E from SARS-CoV-2. The N-terminal region is predicted to be disordered [55]. Representation of the solid-state NMR structure of the SARS-CoV-2 E transmembrane region in DMPC and DMPG model membranes (residues 8-38) [56]. The C-terminal region (residues 41-75) is the focus of this study. **(C)** Sequence alignment of E from SARS-CoV and SARS-CoV-2.

An important question concerns the structural basis for the dual role of E, and for its ability to interact with a variety of different partners. Towards providing an answer to this question, it has been shown that the E-CTD of SARS-CoV exists in a dynamic exchange between different conformations [10]. E-CTD residues 46-75 were identified as a disorder-based protein binding region, also known as MoRF [5], suggesting that the E-CTD may exhibit features characteristic of intrinsically disordered proteins (IDPs), which lack a specific stable three-dimensional structure [11]. The disordered nature of IDPs enables them to adopt different conformations upon binding with different partners [12,13]. In particular, the E-CTD has a predicted β-coil-β motif with a conserved proline residue at its center [10]. Further, the region of residues 46-60 of SARS-CoV E, as a synthetic peptide in isolation, was reported to adopt a β structure under certain conditions [10]. A conserved motif, CxxC (**Figure 1A**), mediates membrane-directed conformational changes [14]. The E-CTD also contains several glycosylation sites, including N45, N48, N4, and N68, which are essential for host membrane-associated oligomerization of E [15]. The last four residues of the E-CTD of SARS-CoV E contain a protein-binding motif known as (PDZ)-binding motif (PBM) (**Figure 1A**), which interacts with host proteins and works to suppress host responses to viral infections [16]. Mutation of the protein binding motif in E-CTD resulted in decreased virulence of SARS-CoV [16,17]. In addition, the short peptide TK9 (TVYVYSRVK) in the SARS-CoV E-CTD, which is important for the oligomerisation of E resulting in the formation of the ion channel complex, has been shown to be amyloidogenic [18].

In this study, in order to provide a more systematic understanding of the conformational properties of the E-CTD, we report a series of measurements under different biological environment mimetics, including in membrane-mimicking environments, under α-helical inducing conditions, and in the presence natural osmolytes (**Figure 1B**). Finally, we investigated the aggregation process of E-CTD upon incubation under physiological conditions.

## Results and Discussion

### The E-CTD is disordered in solution

To obtain insight into the conformational properties of the E-CTD, we performed a sequencebased secondary structure analysis using the s2D, JPred4, PEP2D, and PsiPred servers (**Figure 2**). s2D predicted a tendency to form both ordered and disordered regions in the E-CTD (**Figure 2A**). These results were consistent with those obtained with: (1) JPred4, which predicted residues 6-11 and 16-20 as β-sheet (31%) and remaining random coil (69%) for SARS-CoV-2 (**Figure 2B**), (2) PEP2D, which predicted two regions (residues 7-11 and 20-24) as α-helix, three regions (residues 1-3, 12-16, 27-35) as random coil, and one region (residues 17-19) as β-sheet (**Figure 2C**), and (3) PsiPred, which predicted two regions (residues 41-54 and 64-75) as random coil, one region (residues 55-61) as β-sheet/strand, and one short region (residues 62-63) as α-helix (**Figure 2D**). Taken together, these predictors indicate that the E-CTD has high disordered propensity (55-74%).

**Figure 2.**
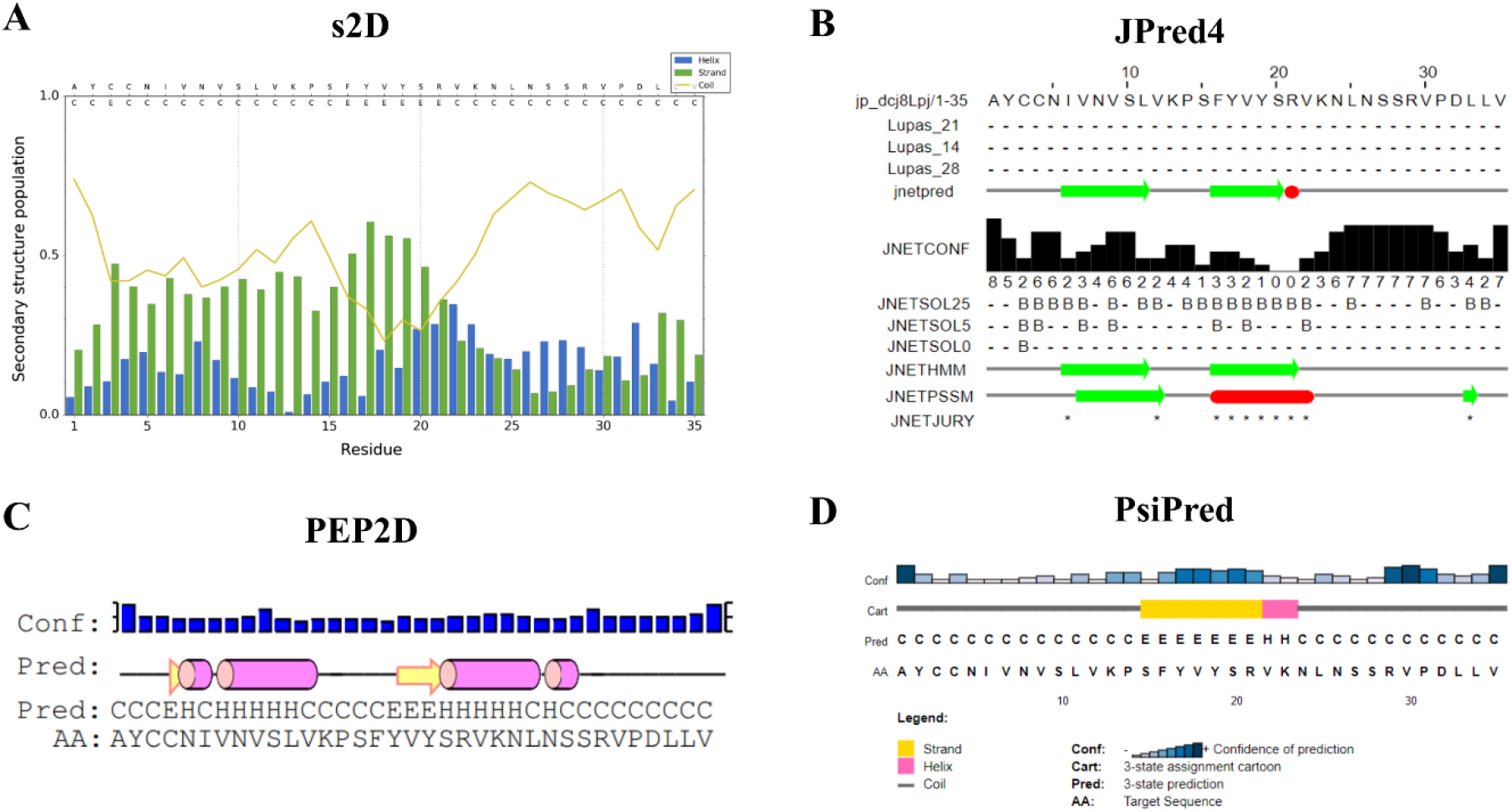
Secondary structure prediction of the E-CTD of SARS-CoV-2 (residues 41-75). **(A-D)** Secondary structure prediction by: **(A)** s2D (blue for α-helices, green for β-strands, and yellow for random coil), **(B)** JPred4, **(C)** PEP2D, and **(D)** PsiPred. H: α-helix, E/B: strand/β-sheet, C: random coil.

Next, using far UV CD spectroscopy, we found that the E-CTD exhibits a minimum at 198 nm (**Figure 3**). The shape of the spectra is characteristic of disordered peptides, consistent with the predictions described above that E-CTD adopts a disordered conformation in an aqueous solution.

**Figure 3.**
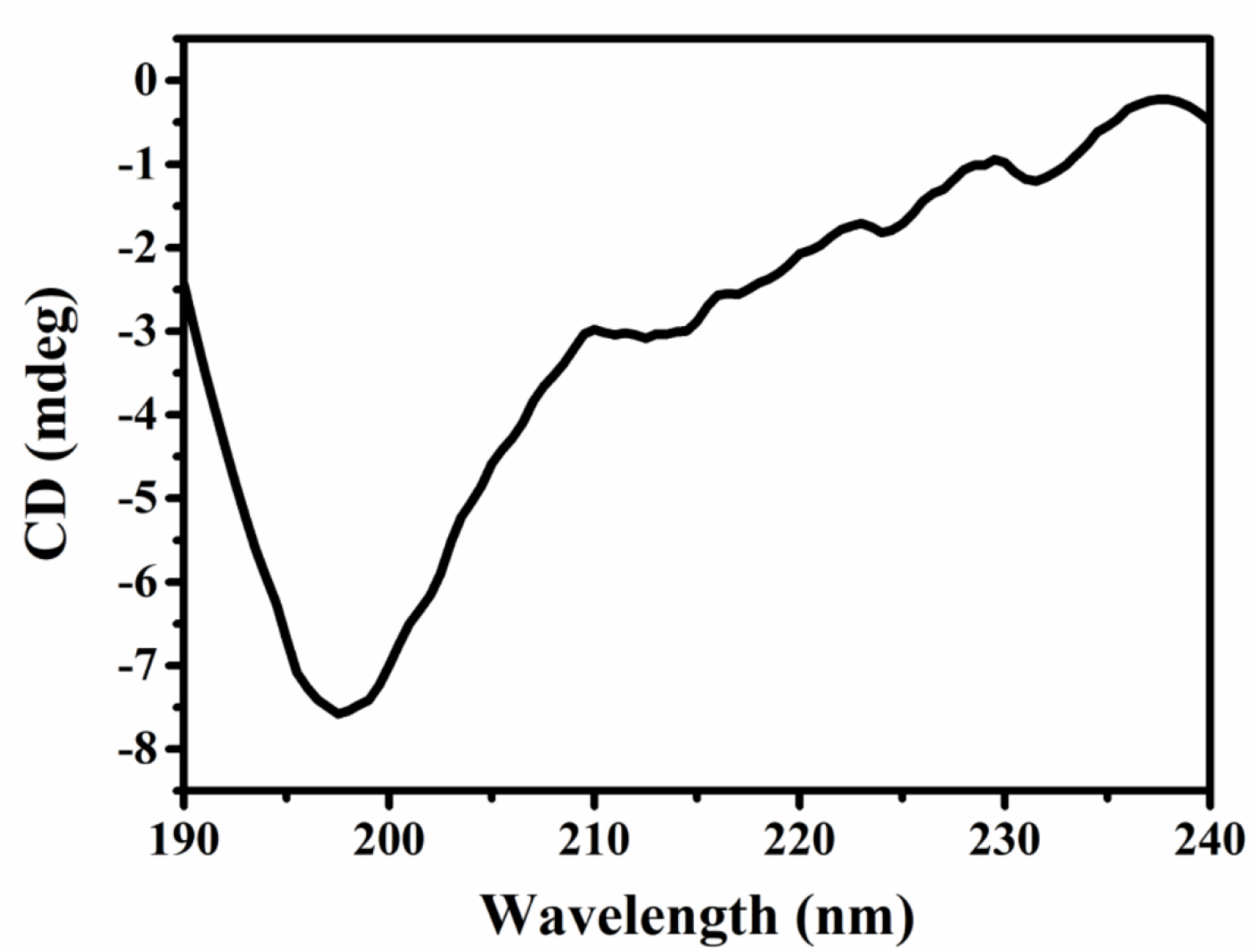
Secondary structure analysis of E-CTD (residues 41-75). Far-UV CD spectrum of 20 μM E-CTD at pH 7.4. The spectrum exhibits a negative peak ellipticity at 198 nm, a signature of disordered proteins.

Further, to obtain insight into the conformational space accessible to the E-CTD, we performed extensive all-atom molecular dynamics simulations up to 1.5 μs. First, we generated an intial structure using homology modelling, finding a structure with α-helices at 43CCNTV47 and 58VYSRVKNLNSS68, while the remaining regions are unstructured. During the simulations, the α-helical content was reduced to 29% in E-CTD, and the unstructured region increased up to 71% (**Figure 4**). The trend for the radius of gyration (Rg) also suggests a dynamic structure, with Rg values varying between 0.9 and 2.2 nm. The large fluctuations in the Rg, and correspondingly in the root mean square deviation (RMSD), is due to the unstructured N-terminal (residues 41-55) and C-terminal (residues 66-75) regions. These results are consistent with the CD spectra described above (**Figure 3**), and with a previous intrinsic disorder analysis of E, which showed a high propensity of the intrinsic disorder [5].

**Figure 4.**
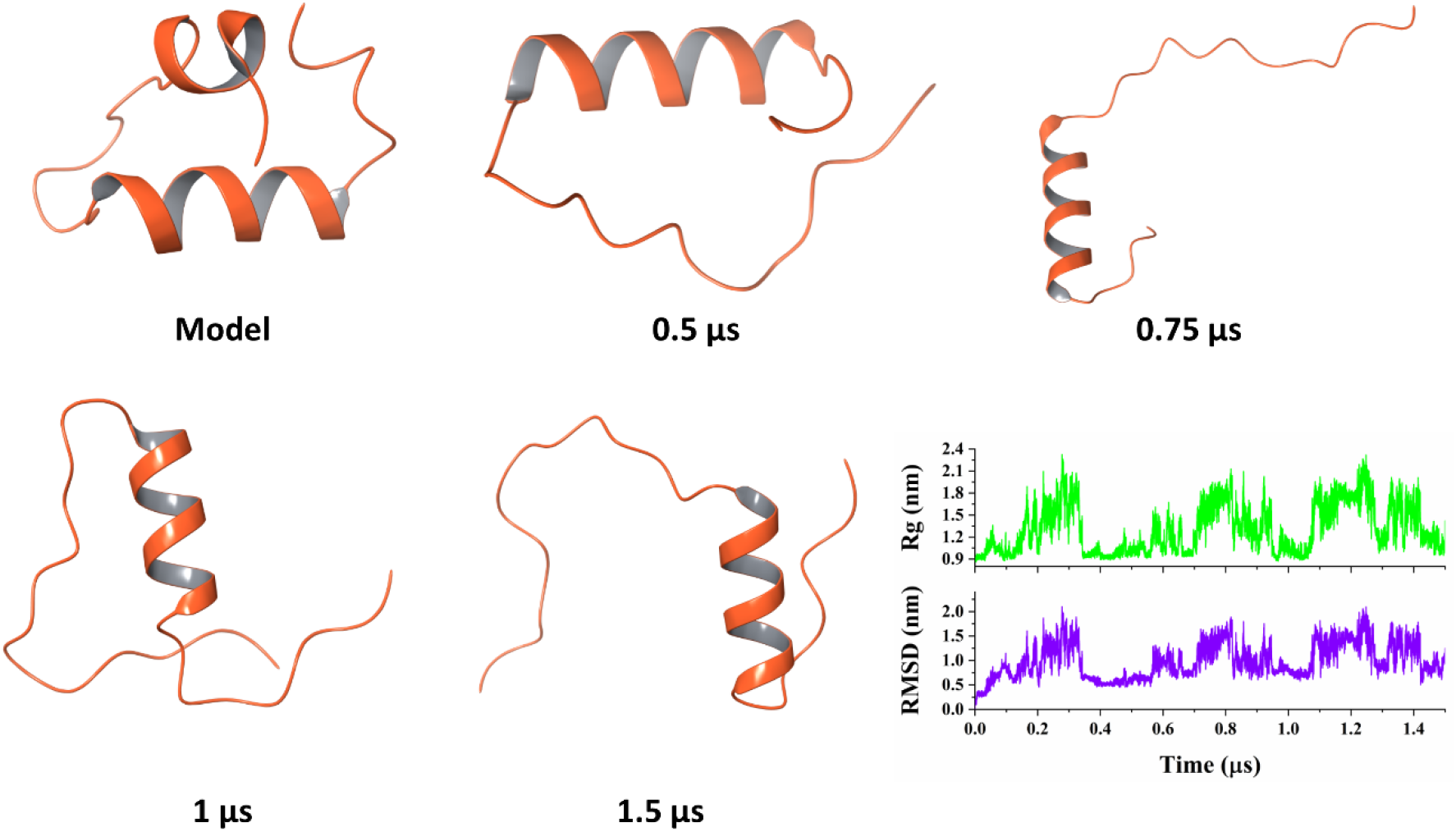
Conformational space explored by the E-CTD (residues 41-75) in MD simulations. Representative conformations were obtained from 1.5 μs MD simulations of the E-CTD in water. The fluctuations of Rg (green) and RMSD (purple) of the E-CTD during the MD trajectory are also shown.

### Conformational properties of the E-CTD in the presence of lipid membranes

It is well known that when certain IDPs approach lipid membranes, they can undergo transient structural changes and form α-helical conformations [19]. Some IDPs can also remain unstructured despite binding with lipid membranes through their side chains [20]. To characterise the conformational properties of the E-CTD in the presence of lipid membranes, we used artificial large unilamellar vesicles (LUVs) to mimic membranous environments. Since E interacts with different cell membranes, we studied how different membrane mimetics, including LUVs (DOPC and DOPS) and SDS micelles, influence the secondary structure of the E-CTD.

We first used DOPC to construct neutral LUVs. Neutral lipids are commonly present at the cell membrane outer leaflet, which is involved in interaction and fusion with a virus [21,22]. Moreover, the Golgi and ER membrane compartments contain a sizeable fraction of neutral lipids [23]. We found that the CD spectra of the E-CTD in the presence of DOPC LUVs exhibited a signature minimum shifted from 198 nm to 218 or 220 nm (**Figure 5A,B**). This result indicates a change in the E-CTD secondary structure, with a gain of β-sheet structure. Since anionic lipids, such as phosphatidylserine, is most commonly found on cytosolic leaflets of Golgi and ER membranes, which are involved in intracellular secretory vesicle trafficking [21,22], we used DOPS to construct anionic LUVs. In this case, the CD spectra of the E-CTD shows a signature minimum at 218 nm, characteristic of proteins with a high content of β-sheet structure (**Figure 5B**). As the concentration of DOPS increased, the signature minimum becomes deeper, suggesting increased β-sheet content. By performing a DichroWeb analysis, we determined the secondary structure content (**Table 3**).

**Table 1.**
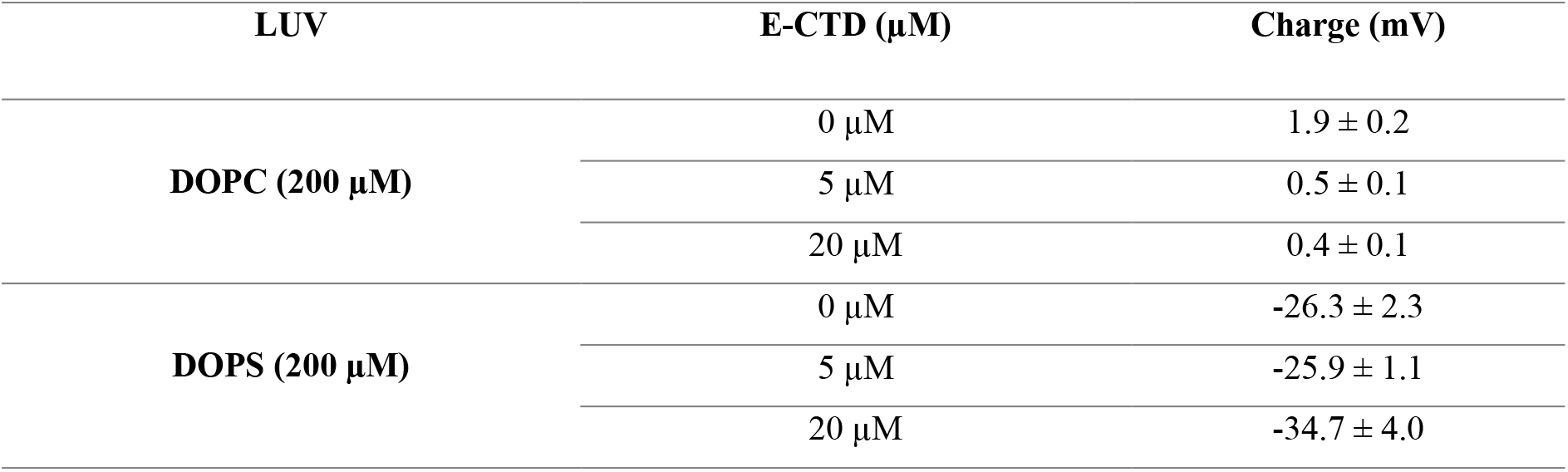
Zeta potential measurement of DOPC and DOPS LUVs in the presence of the E-CTD.

**Table 2:**
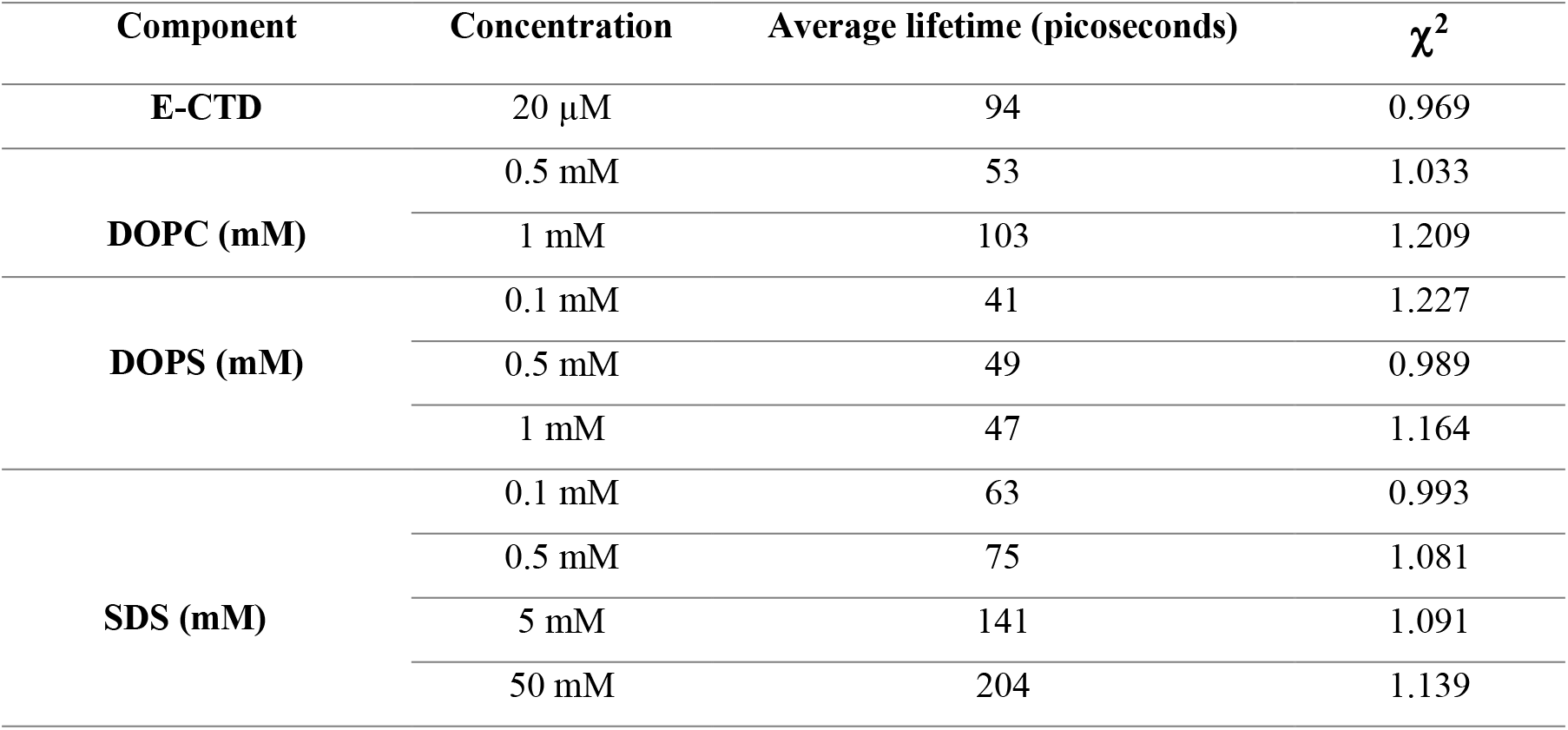
Average lifetime and χ^2^ values for E-CTD in different conditions (DOPC and DOPS LUVs and SDS micelles) after fitting data in three exponential terms.

**Table 3:** Estimation of the secondary structure content of the E-CTD in presence of DOPS LUVs, SDS and TFE using DICHROWEB.

|  |  | Secondary structure content (%) |  |  |  |
| --- | --- | --- | --- | --- | --- |
| | | α-helix | β-sheet | Turns | Disordered |
| **DOPS** | 0 M | 3.8% | 7.0% | 10.6% | 78.5% |
|  | 0.1 M | 8.1% | 14.5% | 23.4% | 53.9% |
|  | 0.5 M | 31.7% | 13.9% | 17.1% | 37.3% |
|  | 1 M | 44.6% | 8.6% | 24.0% | 22.8% |
| **SDS** | 0 mM | 5.45% | 4.3% | 12.7% | 77.5% |
|  | 0.1 mM | 8.9% | 10.4% | 9.1% | 71.5% |
|  | 0.5 mM | 29.8% | 11% | 21.6% | 37.5% |
|  | 5 mM | 39.6% | 17% | 20.1% | 23.2% |
|  | 50 mM | 36% | 8.8% | 12.5% | 42.6% |
| **TFE** | 10% | 2.6% | 6.6% | 10.8% | 79.9% |
|  | 20% | 6.9% | 4.1% | 12.9% | 75.9% |
|  | 30% | 36.2% | 12.5% | 14.95% | 36.3% |
|  | 40% | 42.4% | 12.9% | 16.2% | 28.4% |
|  | 50% | 45.6% | 12.9% | 14.4% | 27% |
|  | 60% | 77.8% | 1.4% | 9.5% | 11.2% |
|  | 70% | 83.4% | 3.6% | 11.8% | 1.1% |
|  | 80% | 82.7% | 3.5% | 9.3% | 4.5% |
|  | 90% | 98.8% | 1.15% | 0% | 0% |

**Figure 5.**
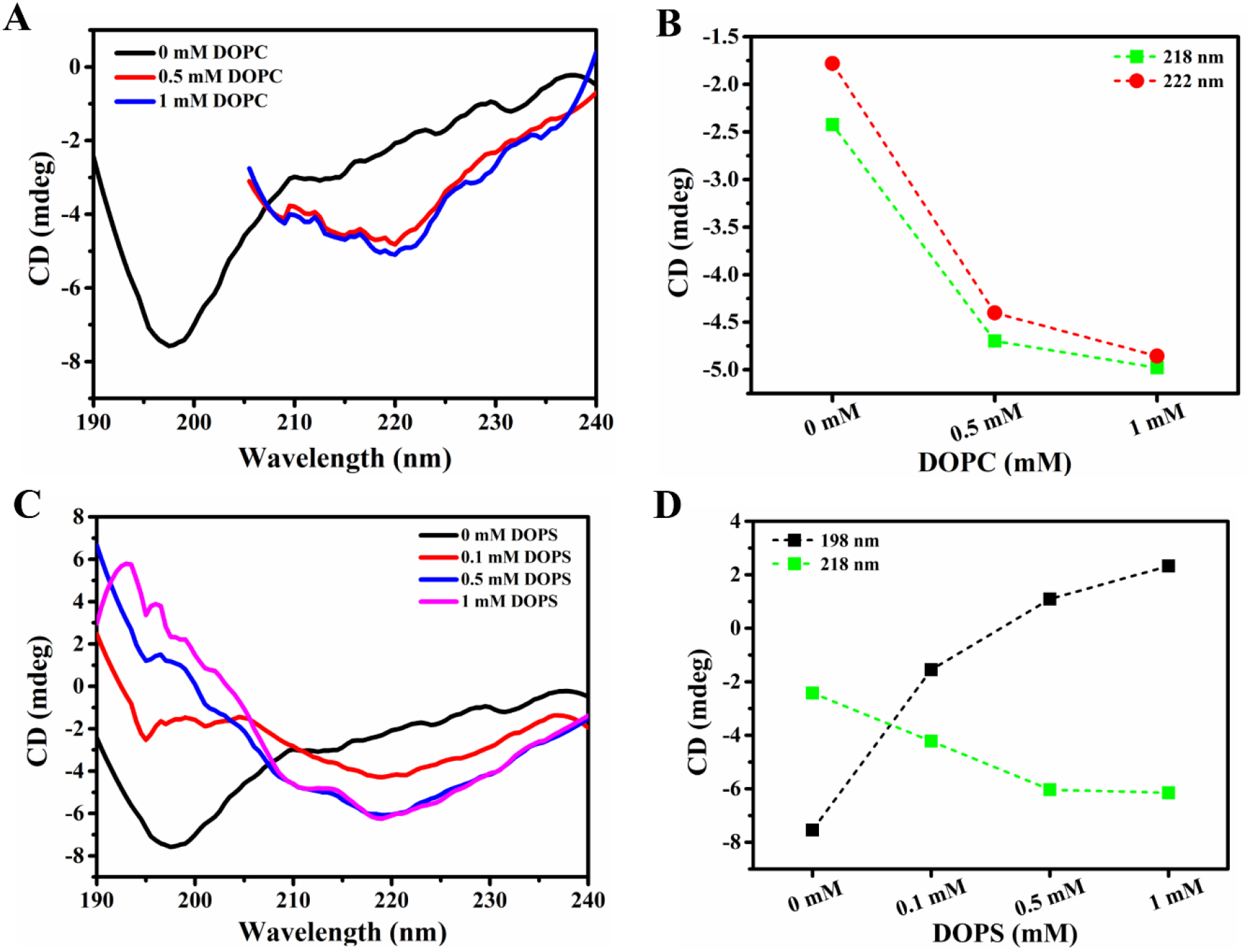
Conformational properties of the E-CTD in the presence of DOPC and DOPS LUVs. **(A)** Far-UV CD spectra of the E-CTD measured at different concentrations of DOPC LUVs, which indicate folding into ordered structures. **(B)** Change of ellipticity at 218 nm (green squares dashed line), and 222 nm (red circles dashed line) on varying DOPC LUVs concentration. **(C)** Far-UV CD spectra of the E-CTD at a varying concentration of anionic lipid DOPS LUVs. **(D)** Change of ellipticity at 198 nm (black squares with dashed line) 218 nm (green squares dashed line) and 222 nm (red circles dashed line) at different concentrations of DOPS LUVs.

Further global conformational dynamics of the E-CTD was assessed by fluorescence spectroscopy and lifetime measurement. Fluorescence spectra show a significant reduction in fluorescence intensity with increasing DOPC concentrations, while the intensity increases with increasing DOPS concentrations (**Figure 6**). These changes in intensity indicate overall conformational changes in the presence of LUVs. The opposite scenarios in the cases of DOPS and DOPC might be due to the total charge on the LUVs, which causes the E-CTD to change the conformation differently.

**Figure 6.**
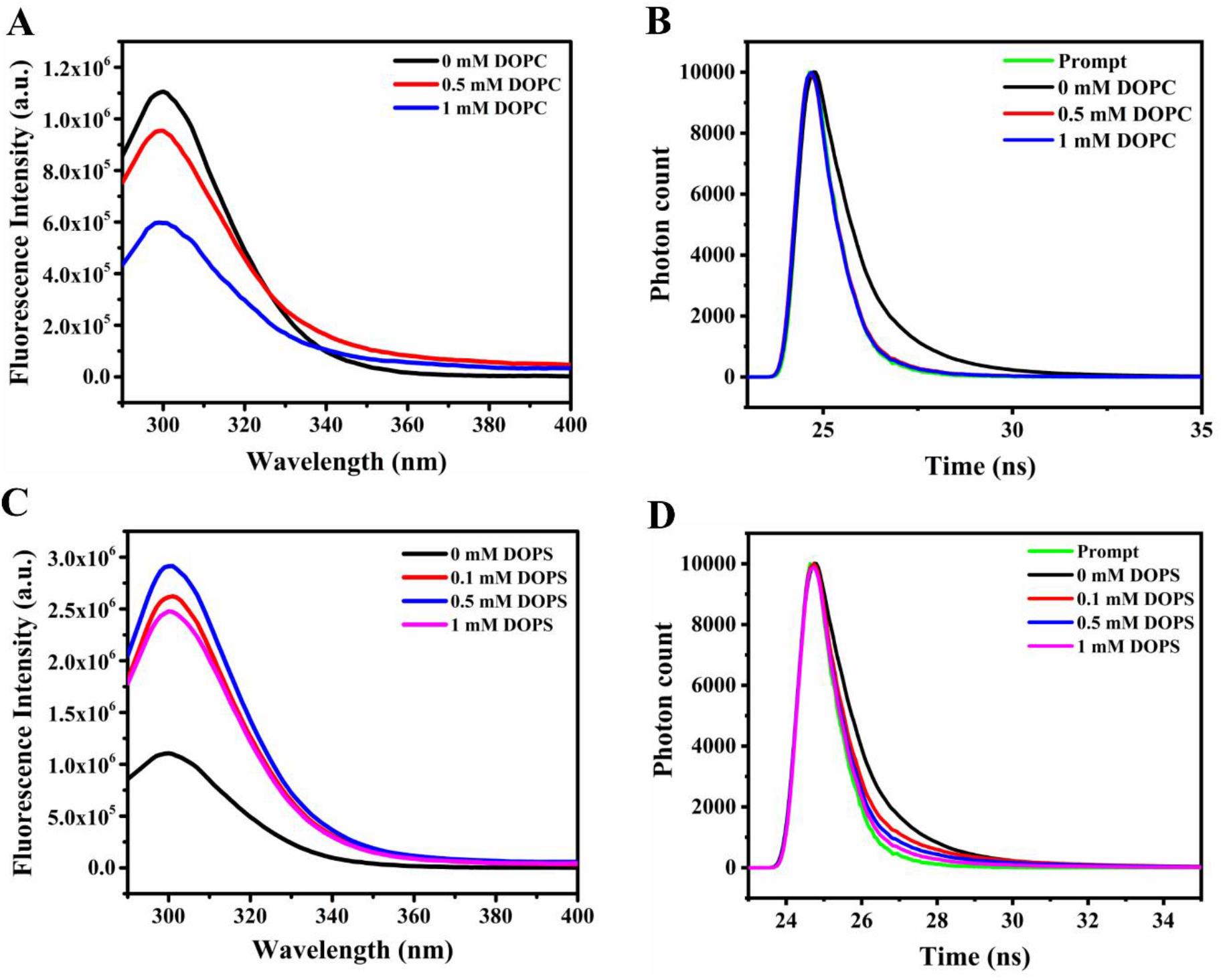
Fluorescence measurements of the tertiary structure of the E-CTD in the presence of DOPC and DOPS LUVs. **(A)** Fluorescence spectra in the presence of DOPC LUVs. **(B)** Fluorescence lifetime decay curve in DOPC LUVs. **(C)** Fluorescence spectra in the presence of DOPS LUVs. **(D)** Fluorescence lifetime decay curve in DOPS LUVs.

Next, to examine the charge of the LUVs by the E-CTD, we measured the zeta potential. We found that the charge of DOPC LUVs was 1.9 ± 0.2 mV, while in presence of 5 μM and 20 μM E-CTD it was slightly reduced to 0.5 ± 0.2 mV and to 0.3 ± 0.1 mV, respectively (**Table 1**). Furthermore, the zeta potential of E-CTD in presence of DOPS LUVs was found to be - 26.3 ± 2.3 mV, while in presence of 5 μM and 20 μM E-CTD, it was −25.9 ± 1.1 mV and −34.7 ± 4.0 mV, respectively (**Table 1**). The analysis of the charge of the LUVs reflects small but significant differences in the presence or absence of the E-CTD. These difference suggest that the E-CTD interaction may perturb the lipid membrane by inducing conformational changes. These changes may occur due to specific inter-molecular interactions of the E-CTD with lipid molecules or salt-bridge interactions, or intra-chain hydrogen bonds [4]. The E-CTD contains the highly conserved cysteine-containing motif CxxC that may allow disulfide isomerization to enhance membrane-directed conformational changes [19]. Moreover, it contains different potent glycosylation sites essential for maintaining the protein broad functional roles and may act as chaperone interacting motifs to help in the protein folding [8,17,41]. The PDZ-binding domain (PBM) motif DLLV at the end of E-CTD plays a crucial role in host cell modifications required for viral infection and pathogenesis [8,24]. The presence of these motifs in the E-CTD may induce membrane-directed conformational changes, which are likely to be relevant in the infectious cycle [37].

### The E-CTD folds in presence of SDS micelles

SDS micelles mimic the interface between hydrophobic and hydrophilic environments, thus enabling the study of protein-membrane interactions [24–26]. SDS is found in micellar form above the critical micellar concentration (CMC) in aqueous conditions (CMC is ~2 mM for SDS micelles in 50 mM phosphate buffer at pH 7), while below the CMC, SDS exists in its monomeric forms [27]. It has been reported that E is dynamic at the membrane interfaces [6]. Here, a substantial change in the shape of CD spectra is seen in the presence of 0.1 mM SDS micelles (**Figure 7A,B**). The difference is prominent at 0.5 mM and above the CMC (5 mM and 50 mM). The spectra in the presence of 0.1-5 mM SDS micelles show a signature minima near 220 nm, which becomes deeper with an increase in SDS micelles, suggesting the gain of β-sheet. Also, in the presence of 0.5 and 5 mM SDS micelles, the spectra show a second minimum near 208 nm, suggesting an α-helix gain. Therefore, at 0.5 and 5 mM, the spectra represent a structure with both β-strand and α-helical propensity. Instead, at higher concentrations of SDS micelles, the spectra exhibit two minima at 222 nm and 208 nm, which indicate the gain of only α-helical structure. Further, the CD deconvolution results by DICHROWEB confirm the decrease in disordered content and an increase in the ordered structural content (**Table 3**) with increasing SDS micelles.

**Figure 7.**
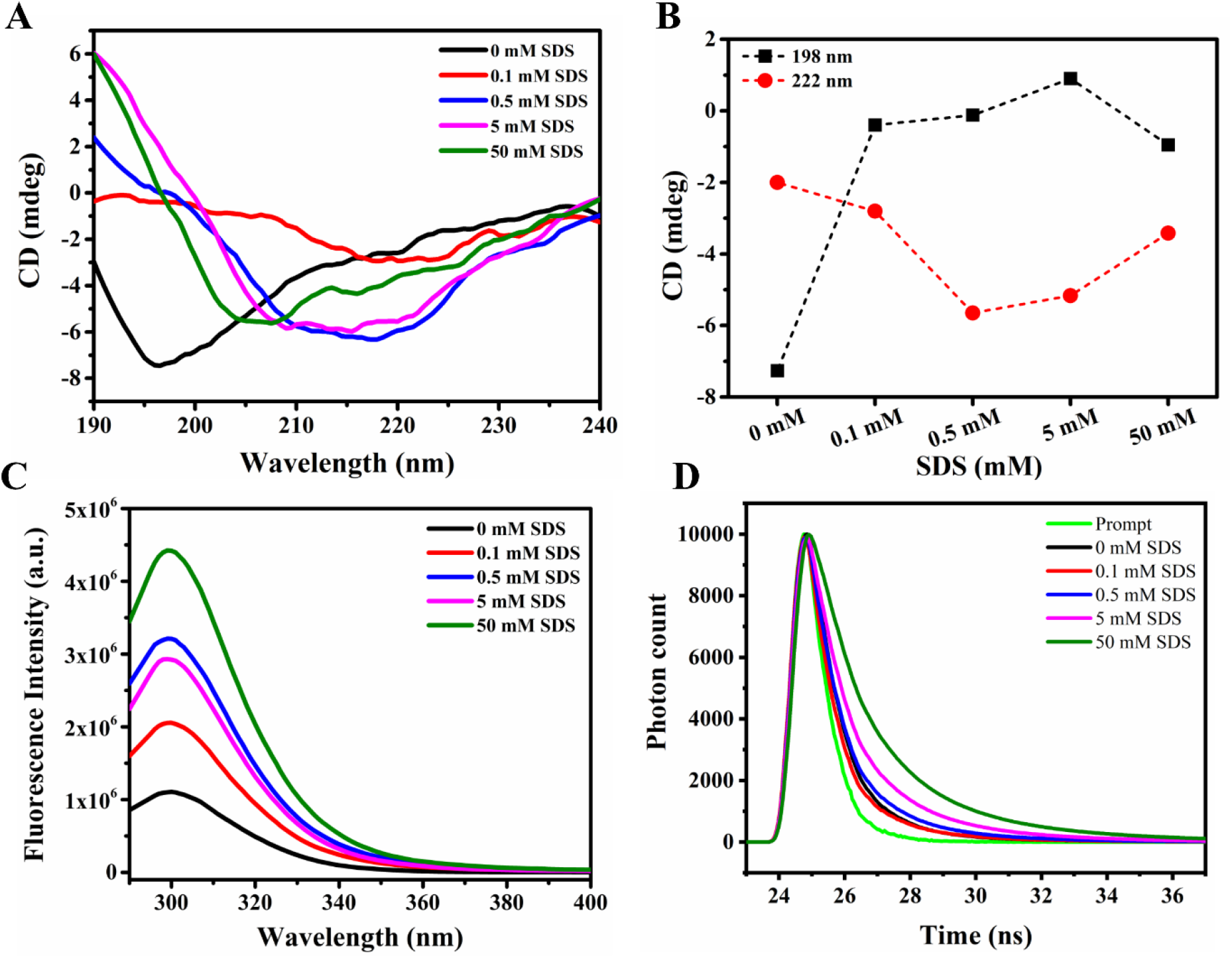
Conformational properties of E-CTD in presence of SDS micelles. **(A)** Far-UV CD spectra in the presence of varying SDS micelle concentration (0-100 mM). **(B)** CD spectra analysis from panel A; ellipticity changes at 198 nm (black squares dashed line) and 222 nm (red circles dashed line). **(C)** Fluorescence emission spectroscopy analysis at different concentrations of SDS micelles. **(D)** Fluorescence lifetime decay curve in varying concentrations of SDS micelles.

Further, the conformational changes observed by fluorescence spectroscopy revealed that with an increasing SDS micelles concentration, the E-CTD shows an increase in fluorescence intensity without a shift in peak position (**Figure 7C,D**). This change in fluorescence intensity may be due to the E-CTD conformational dynamics after interaction with SDS micelles, leading to tyrosine residue exposure. This fluorescence-based result was further confirmed by the lifetime measurement of the tyrosine fluorescence. Any changes in fluorophore physicochemical environments (tyrosine in case of the E-CTD) lead to the change in its property and, consequently, the fluorescence lifetime (**Table 2**). Our findings thus reveal interactions with SDS micelles leading to significant changes in the secondary structure and tertiary structure of E-CTD.

### TFE induces a-helical structure in the E-CTD

Further, to explore the disorder to order transition in the E-CTD, we used trifluoroethanol (TFE), an organic solvent known to induce α-helical secondary structure in proteins. The amphiphilic TFE molecules surround a protein, weakening the hydrogen bonding of water to the CO and NH groups in the coil form [28]. This process leads to dehydration of the protein backbone, which causes the formation of backbone-backbone hydrogen bonds and therefore induces the formation of α-helical structure [29–31]. Other studies also proposed that TFE act to destabilize the unfolded state by structuring the solvent, leading to an elevated folded population [32,33]. We measured CD spectra of the E-CTD in varying concentrations of TFE (**Figure 8A**). As TFE concentration increases, the signature minimum at 198 nm is progressively lost, and new signature minima appear at 208 nm and 222 nm (**Figure 8B**). These results indicate that the E-CTD acquires an α-helical structure in the presence of TFE. A percentage secondary structure analysis by DICHROWEB shows a gain of α-helix content to 99% at 90% TFE (**Table 3**). Further, fluorescence spectroscopy demonstrated that the E-CTD displays an increase in intensity upon increasing TFE concentrations (**Figure 8C**).

**Figure 8.**
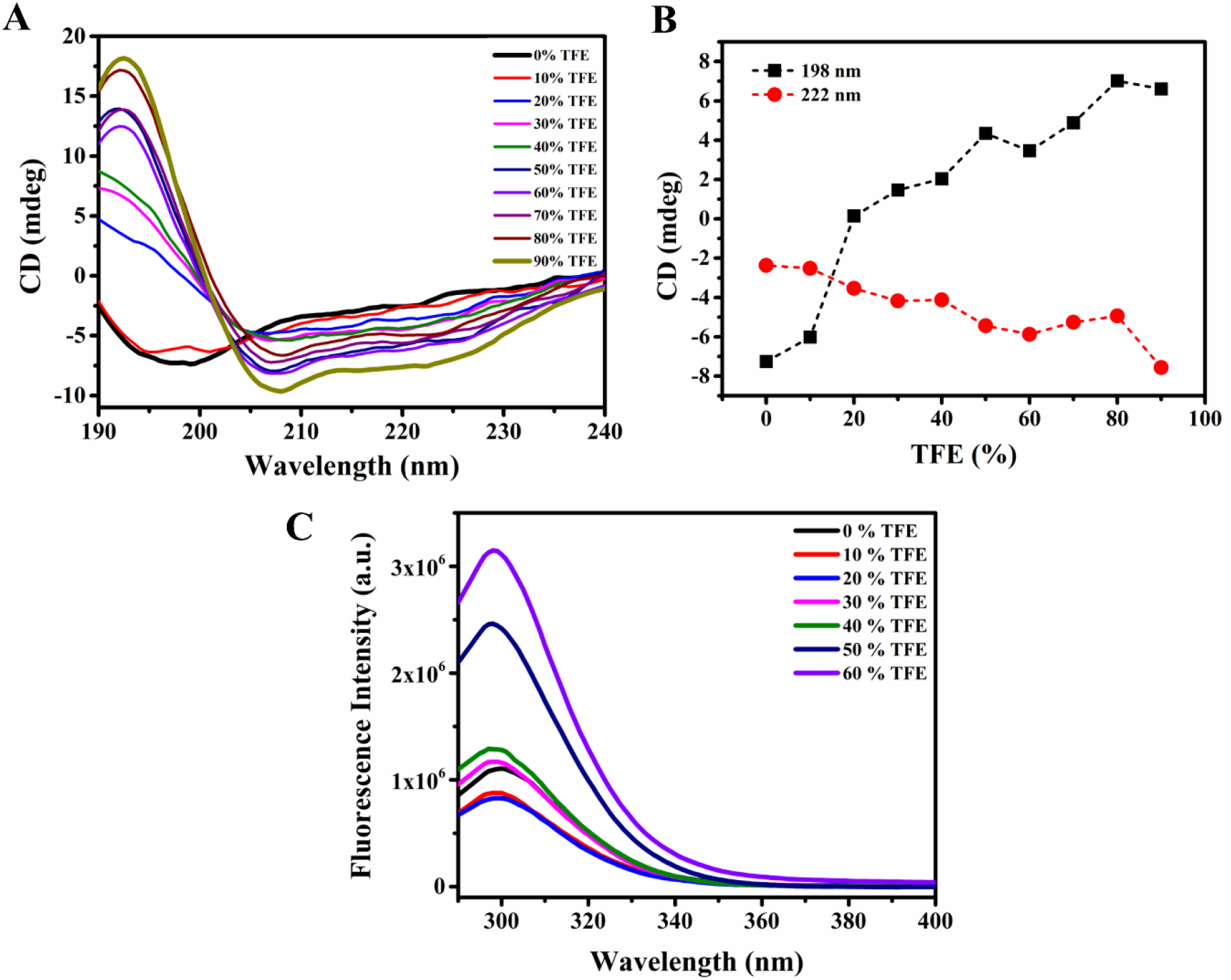
TFE induces α-helical structure in the E-CTD. **(A)** Far-UV CD spectra in the presence of different concentrations of TFE. **(B)** Analysis of CD spectra from panel A. Ellipticity change at 198 nm (black squares dashed line) and 222 nm (red circles dashed line). **(C)** Fluorescence emission spectroscopy analysis.

### Natural osmolytes enhance the folding of the E-CTD

Osmolytes are naturally occurring low molecular weight organic molecules that help evade stress conditions and regulate several biological processes such as protein folding, protein-protein interactions, and protein disaggregation [34–36]. Since stress conditions could follow SARS-CoV-2 infection, we investigated the effects of osmolytes on the conformational properties of the E-CTD. The far-UV CD spectra of E-CTD in the presence of trimethylamine N-oxide (TMAO), an organic compound that stabilizes structured states of proteins, show an increased β-strand content, which is evident from the observed CD signal at 218 nm (**Figure 9A**). In the presence of TMAO, the E-CTD exhibits a change in the structural conformation into an ordered form. We also observed a transition from 0 M TMAO to 2.5 M TMAO. At 0.2 M TMAO, the E-CTD shows reduced negative ellipticity but does not show a significant change in ellipticity at 222 nm (**Figure 9B**). At 1 M and above, a gradual increase in ellipticity at 222 nm indicates increased β-strand content. Disordered proteins have been shown to fold in the presence of TMAO [37–39]. However, some IDPs exhibit a different behaviour [40,41], as they only exhibit a reduction in negative ellipticity at 198-202 nm. Overall, these results indicate that the E-CTD in the presence of TMAO undergoes a transition from disorder to ordered (**Figure 9**). Moreover, these structural changes were further confirmed by intrinsic fluorescence experiments that show a reduction in fluorescence intensity with increasing TMAO concentration.

**Figure 9.**
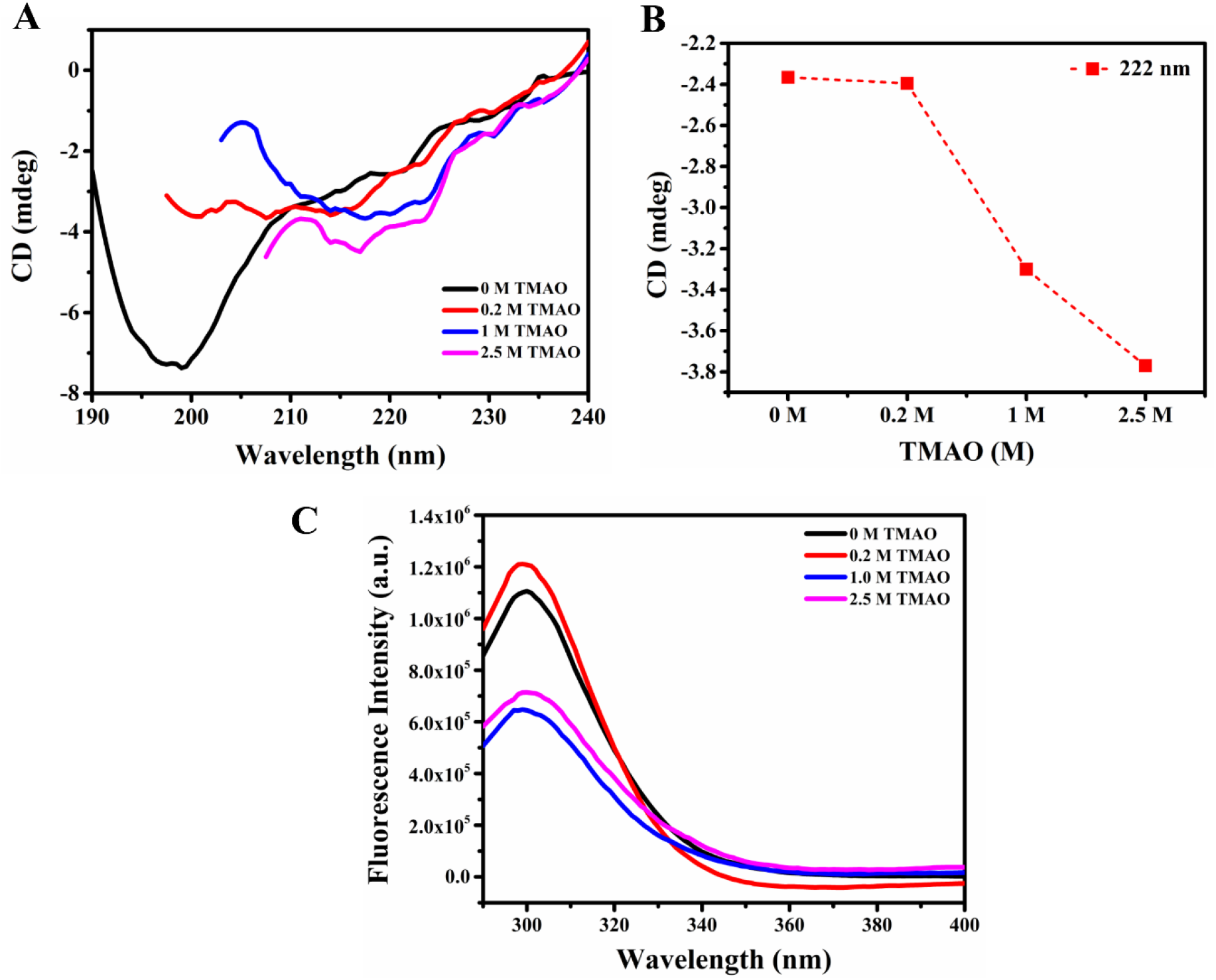
Conformational properties of the E-CTD in presence of TMAO. **(A)** Far-UV CD spectra in the presence of different concentrations of TMAO. **(B)** Analysis of the CD spectra from panel A at 222 nm (red circles with dashed line). **(C)** Fluorescence emission spectroscopy analysis.

### The E-CTD shows a temperature-dependent compaction characteristic of IDPs

Temperature-induced structural changes are common among IDPs, including a conformational ensemble compaction [11,42]. To investigate the temperature-induced structural changes in the E-CTD, we recorded far-UV CD spectra at increasing temperatures (10 °C to 90 °C, at 5 °C intervals). We observed typical CD spectra for disordered proteins at low temperatures. However, upon increasing the temperature, the shape of the CD spectra changed (**Figure 10A**). A significant gain in negative ellipticity nearby 222 nm, and a loss of negative ellipticity around 198 nm, were observed (**Figure 10B**). This type of temperature-driven structural change is known to derive from structural compaction, as observed in other disordered proteins and peptides [39,41,43–47]. However, other studies on IDPs showed that disordered proteins may lose their α-helical structure with increasing temperature, suggesting that the α-helices are not solely responsible for the spectroscopic change detected by the CD spectroscopy [11]. Thus, the temperature-induced change in the E-CTD may be interpreted as heating-induced folding of the E-CTD into α-helical structures.

**Figure 10.**
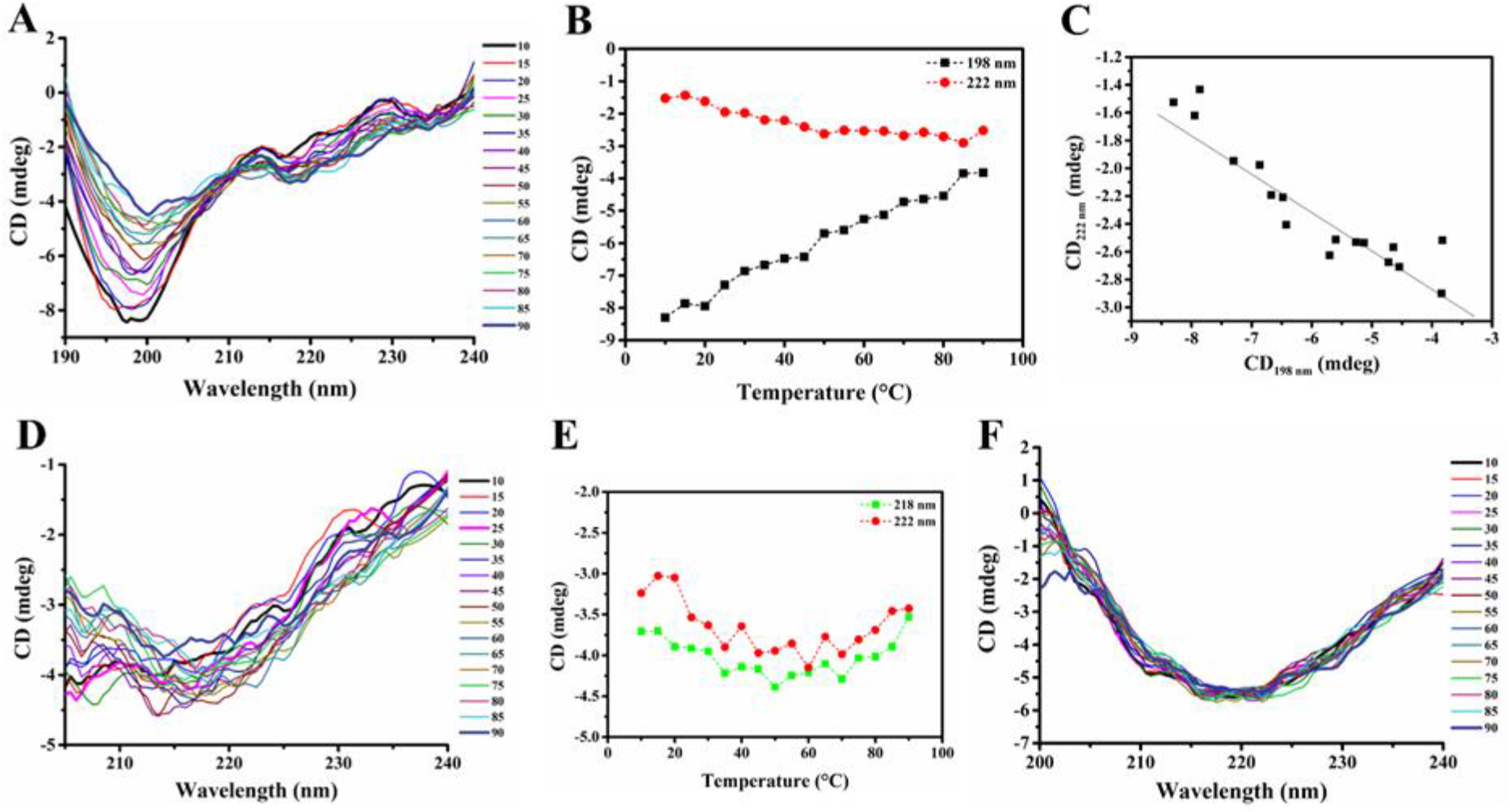
Effects of temperature on the secondary structure of the E-CTD. (**A**) The thermal stability at varying temperature conditions was monitored using CD spectroscopy at 10 °C to 90 °C, with 5 °C intervals. (**B**) Analysis of spectral changes by examining ellipticity changes at two wavelengths as 198 nm (black line) and 222 nm (red line). (**C**) Phase diagram obtained by plotting CD data at 198 nm (x-axis) and 222 nm (y-axis). (**D**) Structural analysis of the E-CTD in the presence of DOPC LUVs at varying temperatures. (**E**) Analysis of spectral changes for the data in panel D by examining ellipticity change at two wavelengths as 218 nm (green line) and 222 nm (red line). (**F**) Structural analysis of the E-CTD in the presence of DOPS LUVs at varying temperatures.

Furthermore, to study the influence of temperature on the secondary structure of the E-CTD in the presence of lipid membranes (DOPC and DOPS LUVs), we recorded CD spectra with varying temperatures (**Figure 10**). At room temperature, DOPC and DOPS LUVs induce ordered secondary structure in the E-CTD (**Figures 5** and **6**), and the shape of spectra remains the same irrespective of temperature increase. The presence of lipids stabilizes the β-sheet in the E-CTD even at higher temperatures confirming the interaction between the E-CTD and lipids.

### The E-CTD forms amyloid-like fibrils

The fluorescent dyes ThT and ANS show strong binding to E-CTD aggregates (**Figure 11A, B**). ANS shows a significant blue shift upon binding E-CTD aggregates compared with controls and freshly dissolved E-CTD. It also represents secondary structural changes (**Figure 11C**), a reduction in spectral ellipticity were observed along with a shift of spectra to gain ordered structure. Using AFM, we confirmed the presence of amyloid-like fibrils after incubation for 45 days of the E-CTD under physiological conditions (**Figure 11D**), with a height of approximately 1.3 nm (**Figure 11F**).

**Figure 11.**
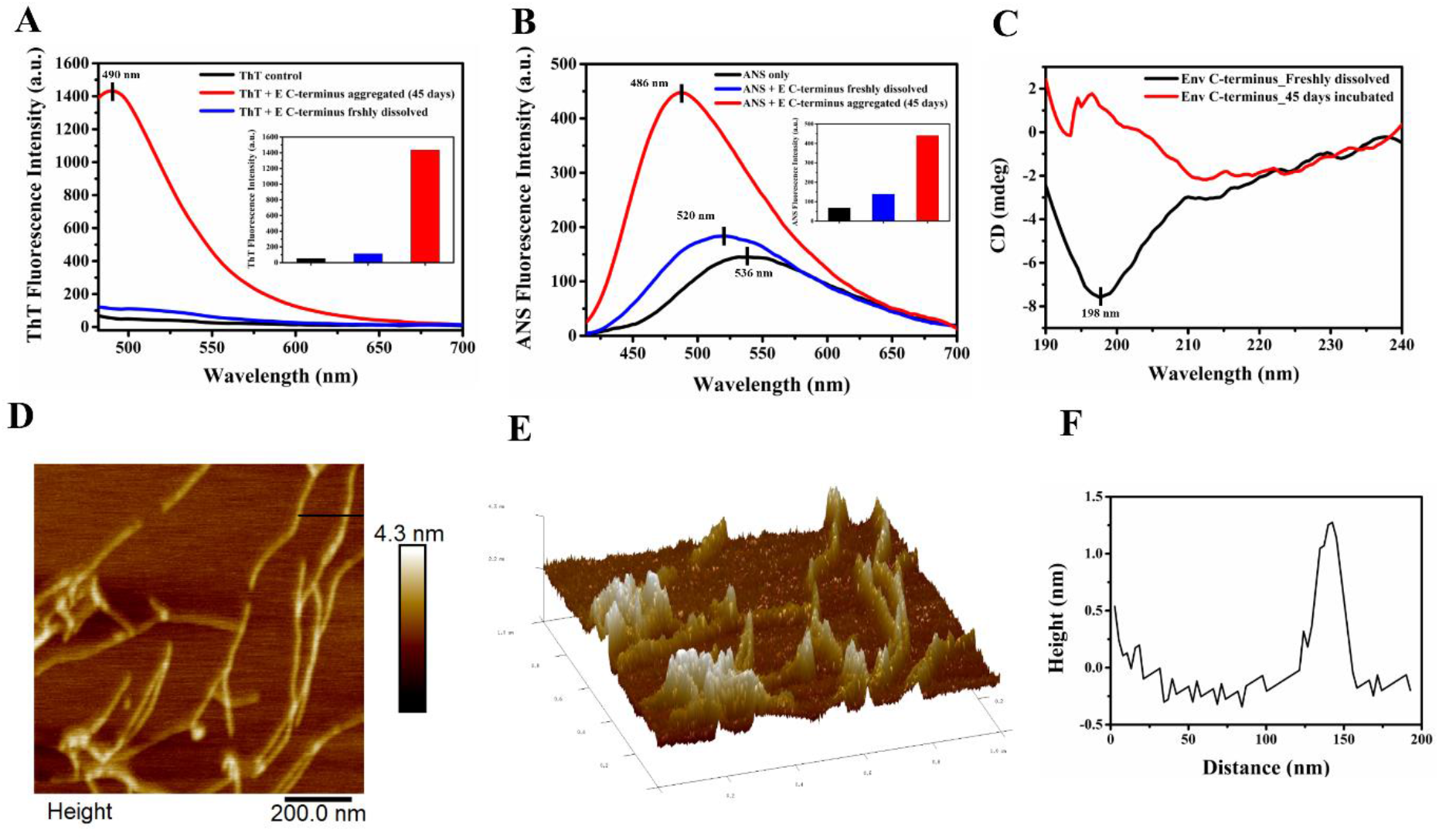
Amyloid formation by E-CTD. (**A**) ThT fluorescence emission spectra from 470 to 700 nm at 450 nm excitation wavelength. (**B**) ANS fluorescence. (**C**) CD spectra; the black line represents freshly dissolved E-CTD, and the red line represents aggregated E-CTD (45 days). (**D**) AFM image for a sample incubated for 45 days; the scale bar indicates 200 nm. (**E**) 3D topography image of E-CTD fibrils from an image of panel D. (**F**) Height line profile of E-CTD fibrils from an image of panel D.

## Conclusions

We have investigated the structural basis for the interaction of the SARS-CoV-2 envelope protein (E) with host cell membranes. We have shown that the C-terminal domain of E (E-CTD) in solution is intrinsically disordered, but also that it changes dramatically its structural behavior in the presence of membrane mimetics, osmolytes, and organic solvents. We have observed that the E-CTD exhibits a structural transition from disorder to order upon interacting with LUVs, SDS micelles, TMAO and TFE. The E-CTD adopts an α-helical structure in TFE, a β-sheet structure in DOPS and DOPC LUVs, indicating that lipid membranes may influence its secondary structure. These conformational changes may have an important role in the viral infection cycle. In addition, upon incubation at physiological conditions, the E-CTD formed amyloid-like fibrils.

These results indicate that the E-CTD can respond to changes of environmental conditions by adopting different conformations, a dynamic behavior that may be critical for viral infectivity. We anticipate that to fully elucidate the heterogeneous nature of the SARS-CoV-2 E-CTD, further studies will be needed to link mechanistically the conformational dynamics in viral pathogenesis.

## Experimental procedures

### Materials

The C-terminal sequence of the envelope protein (residues 41–75, [NH2]-AYCCNIVNVSLVKPSFYVYSRVKNLNSSRVPDLLV-[COOH]) was obtained from Genscript USA Inc., with >76.9% purity. Sodium dodecyl sulfate (SDS), trimethylamine N-oxide (TMAO), 2,2,2-trifluoroethanol (TFE), thioflavin T (ThT), and 8-anilinonaphthalene-1-sulfonic acid (ANS) were purchased from Sigma-Aldrich (St. Louis, USA). 1,2-dioleoyl-sn-glycero-3-phospho-L-serine (DOPS) and 1,2-dioleoyl-sn-glycero-3-phosphocholine (DOPC)) were purchased from Avanti Polar Lipids (Alabaster, Alabama, U.S.A.).

### LUVs preparation

Lipids (DOPC and DOPS) were dissolved in chloroform, and the chloroform was removed from the solution using a rotatory evaporator. Further, dried lipids were hydrated with 50 mM sodium phosphate buffer (pH 7.4), and the lipid suspension (the concentration of DOPC suspension was 29 mg/ml, and the concentration of DOPS suspension was 20 mg/ml) was processed five times by freeze (liquid N2)/thaw (60 °C water bath)/vortex cycles. The lipid suspension was then subjected to extrusion (Avanti Mini Extruder) to prepare LUVs using 0.1 μM pore diameter polycarbonate membrane. The prepared LUVs were stored at 4 °C and used within seven days. Detailed protocols have been reported previously [39,41,47].

### Dynamic light scattering (DLS)

LUVs composed of DOPC or DOPS were diluted in water (1:100 ratio of LUV and water), and their size was measured by using Malvern Zetasizer Nano S (Malvern Instruments Ltd., UK). The observed size of the DOPC and DOPS LUVs was 102 nm and 124 nm, respectively.

### Circular dichroism (CD) spectroscopy

The CD spectra of the E-CTD were recorded on JASCO (J-1500) spectrophotometer furnished with the temperature regulator and Peltier cell holder. Far-UV CD spectra were recorded as an average of 3 consecutive scans. 1 mm optical path length quartz cuvette was used to record CD spectra. A 20 mM phosphate buffer at pH 7.4 was used to prepare the E-CTD sample at 20 μM. CD spectra were recorded for the E-CTD alone at 20 μM and in the presence of various concentrations of DOPC and DOPS LUVs, SDS, TFE, and TMAO at 25 °C. Each spectrum was subtracted from that of the buffer blank. Further, the smoothing of CD spectra was done by Savitsky-Golay fitting at 15 points of smoothing window and second polynomial order. Finally, secondary structure contents were calculated by deconvoluting the CD spectra in the CONTIN algorithm with a reference set 7 using the DICHROWEB web server [48].

### Secondary structure prediction

We used different web servers, including s2D, PSIPRED, JPred and PEP2D, to predict the consensus secondary structure of the E-CTD of SARS-CoV and SARS-CoV-2.

### Fluorescence measurement

Since E-CTD has three tyrosine (Tyr) residues, Tyr was used as an intrinsic fluorophore to monitor the fluorescence emission intensity using a Fluorolog spectrofluorometer (HORIBA Scientific) with an excitation wavelength of 260 nm. A 20 μM protein sample was prepared in 50 mM sodium phosphate buffer at pH 7.4, and monitored fluorescence emission in 5 nm slit bandwidth in a quartz cuvette (400 μl sample volume).

### Fluorescence lifetime measurement

The fluorescence lifetime of Tyr in the E-CTD was measured using the DeltaFlex TCSPC system (Horiba Scientific). The wavelengths for excitation monochromator was set up at 280 nm and emission monochromator at 310 nm. The measurement range was set up to 200 ns with 32 nm of bandpass and a peak preset of 10000 counts. Ludox was used to correct the instrument response factor (IRF), and the wavelength for prompt measurement was set up at 280 nm. 20 μM protein sample was prepared in 50 mM sodium phosphate buffer at pH 7.4.

### Molecular dynamic (MD) simulations

As no structure is currently available for the E-CTD, initial models were generated using the PepFold web server using 200 simulations of energy minimization. The starting model for the MD simulations was built and then prepared using Chimera by employing a previously described protocol [41,47]. To examine the conformational space accessible to the E-CTD, we performed all-atom MD simulations up to 1.5 μs using the Charmm36 force field [49] in Gromacs v5 [50], as reported previously [51]. In the presence of TIP3P water model, 0.15M NaCl, and solvation neutralizing counterions, the structure of the E-CTD was prepared and minimized using steepest descent method. With NVT and NPT equilibration at 300 K and 1 bar pressure, the system was equilibrated. Using Nose-Hoover and Parrinello-Rahman coupling methods, the average temperature at 300 K and pressure at 1 bar were maintained during the simulations. The final production run was performed for 1.5 μs using the high performing cluster at IIT Mandi.

### Thioflavin T (ThT) binding assay

The benzothiazole dye thioflavin T (ThT) is a frequently used probe for in vitro amyloid fibril formation by proteins/peptides [52]. ThT forms highly fluorescent complexes with amyloid fibrils and does not interact with the globular proteins in a native state [53]. In this study, we used ThT to identify fibril formation by the E-CTD. 20 μM ThT was mixed with 50 μM E-CTD samples (monomeric and aggregated) in sodium phosphate buffer (20 mM phosphate, 50 mM NaCl, pH 7.4) and incubated for five minutes in the dark. Emission spectra were recorded in triplicate at 450 nm excitation wavelength and emission scan from 470 nm to 700 nm in a TECAN Infinite M200 PRO microplate reader. The final spectra represent the average of three spectra.

### ANS fluorescence assay

The ANS dye was used to detect partially folded intermediates such as molten globules and measuring the exposed hydrophobic surface during amyloid fibril formation [54]. E-CTD samples (fresh and incubated ofr 45 days) were mixed with ANS (in 20 mM sodium phosphate buffer, pH 7.4) at a final concentration of 45 μM, respectively. Emission spectra were recorded from 400 nm to 700 nm with 380 nm excitation wavelength in TECAN Infinite M200 PRO multimode microplate reader. The spectra were acquired in triplicates, and the final spectra represent the average of three measurements.

### Atomic force microscopy (AFM)

AFM images of E-CTD fibrils were acquired in tapping-mode AFM (Dimension Icon from Bruker). Aggregated samples were diluted thirty times with deionized water and mounted on a freshly cleaved mica surface and incubated for 1-hour. Further, samples were rinsed with deionized water, dried overnight, and images were recorded.

## Data availability

All the data are contained within the manuscript.

## Authors Contribution

RG, NG, and MV: Conception, design, review, and writing of the manuscript. KG and AK performed experiments, analyzed data, and wrote the manuscript. PK performed all the computational simulations, analyzed, and wrote the respective part. SKK helped in figure preparation and writing the manuscript.

## Acknowledgements

All authors would like to thank Indian Institute of Technology Mandi (Bio-X center, HPC facility, AMRC center & C4FED Clean Room) for all the facilities and faculty research grant, School of Basic Sciences (SBS), IIT Mandi. KG was supported by the fellowship from the Science and Engineering Research Board (SERB), India (Grant Number: CRG/2019/005603). AK is thankful to Ministry of Human Resource Development (MHRD) for fellowship. SKK were supported by the Indian Council of Medical Research (ICMR), India, for fellowship. RG is grateful to IYBA Award (Grant Number: BT/11/IYBA/2018/06) by Department of Biotechnology (DBT), India, and MHRD-SPARC (SPARC/2018-2019/P37/SL).

